# *De novo* assembly and annotation of the *Empoasca fabae* mitochondrial genome

**DOI:** 10.1101/2024.09.20.614148

**Authors:** Joshua Molligan, Jordanne Jacques, Soham Mukhopadhyay, Edel Perez-Lopez

**Affiliations:** Département de phytologie, Faculté des sciences de l’agriculture et de l’alimentation, Université Laval, Québec City Québec, Canada; Centre de recherche et d’innovation sur les végétaux, Université Laval, Québec City Québec, Canada; Institute de Biologie Intégrative et des Systèmes, Université Laval, Québec City Québec, Canada; L’Institute EDS, Université Laval, Quebec City, Québec City, Québec, Canada

**Keywords:** Cicadellidae, mitogenome, *Empoasca* sp., leafhoppers genomics, next generation sequencing

## Abstract

This study presents the assembly and annotation of the full-length mitochondrial genome for the leafhopper species *Empoasca fabae* Harris, 1841. The mitogenome was obtained from a contig-level assembly with the identified mitochondrial genome being 14,873 bp in length. The base composition was A (38.8%), T (39.1%), C (11.7%), and G (10.4%). The mitogenome comprised 13 protein-coding genes (PCGs), 22 transfer RNA genes (tRNAs), two ribosomal RNA genes (rRNAs), and showed a unique, non-AT-rich D-loop region. Nearly all PCGs being with an ATN start codon, while two begin with TCG and GTG. Phylogenetic analysis confirmed the placement of *E. fabae* within the subfamily Typhlocybinae, clustering with other species in the *Empoasca* genus.

## INTRODUCTION

*Empoasca fabae* Harris, 1841, is a significant insect pest in North American agriculture, belonging to the family Cicadellidae in the order Hemiptera. Its economic impact stems from its polyphagous feeding habits, annual migratory patterns, and suspected ability to transmit viral and bacterial plant pathogens across a wide range of economically important crops (Santos Almeida et al. 2024a, 2024b). Our group has also been using this migratory leafhopper as a model to study the effects of climate change on insect migration, population dynamics, disease transmission, and insecticide resistance (Plante et al. 2024). To advance this research, population genomic studies are essential. However, we are currently facing a bottleneck due to the lack of available mitochondrial genomes for *E. fabae*, which has prompted us to produce and annotate the first complete mitochondrial genome of this species using high-throughput Illumina sequencing.

## MATERIALS AND METHODS

### Sample Collection and DNA Extraction

Leafhoppers used in this study were captured using yellow sticky traps between July and August 2023 from a strawberry farm in Southern Québec, Canada (45°34’24.0”N 73°03’46.0”W). Samples were visually identified using taxonomic keys to the species level as previously described (Plante et al. 2024), preserved in 70% ethanol and stored at 4°C until DNA extraction.

*E. fabae* specimens were pooled into 3 subsamples of 10 insects each and washed with sterile ddH□O. DNA was then extracted by homogenizing the insects with a mini-pestle in 700µL of lysis buffer containing 20mM EDTA (pH 8.0), 100mM Tris-HCl (pH 7.5), 1.4M NaCl, 2% w/v CTAB, and 4% w/v PVP-40. After homogenization, an additional 700µL of lysis buffer was added, and samples were incubated at 65°C for 1 hour, inverting the tubes every 10 minutes. The supernatant was then washed twice with chloroform-isoamyl alcohol (24:1), and DNA was precipitated using 70% v/v ice-cold isopropanol. The DNA pellet was washed with 70% ethanol and air-dried for 10 minutes before eluting in a buffer containing 10mM Tris-HCl and 0.1mM EDTA at a pH of 8.0.

### Sequencing and Preprocessing

The quality of DNA was evaluated using Nanodrop (Thermo Fisher Scientific, USA) and Qubit (Thermo Fisher Scientific, USA). Preparation of Illumina short-read libraries and subsequent sequencing were performed by Genome Québec (Montréal, Canada), resulting in three paired-end (PE) files of 43M, 49M, and 50M reads, respectively. Raw data files were preprocessed with BBTools (v.36.92) to trim adapter sequences (k=23, mink=11, hdist=1), with flags for paired-end trimming and overlap detection (Bushnell, 2024). Additional quality filtering was performed (trimq=10), and low complexity sequences were removed (entropy=0.7, entropywindow=50, entropyk=5). Finally, reads shorter than 20bp were then filtered and discarded.

### Assembly and Mitogenome Identification

After preprocessing, the forward and reverse PE files were merged, totaling 142M reads. A *de novo* assembly was performed using SPAdes (v.4.0.0), with an adjusted kmer size (21, 33, 55, 77, 99, 127) (Prjibelski et al., 2020). The assembly used repeat resolution, mismatch careful mode, and a mismatch corrector to polish contigs. This produced a total of 2.5M contigs, which were filtered for contigs ≥ 5kb and ≥ 5x coverage, reducing the total to 1,583 contigs.

The filtered contigs were blasted using NCBI BLAST (v.2.13.0) to identify top 10 matches with > 80% identity (Altschul et al., 1990). Contigs were further filtered for > 40% coverage to the queried nodes. The top matches were retrieved, and their accession numbers were then queried with edirect (v.14.6) to retrieve taxonomy data from the nucleotide database (NCBI, 2024). Out of 41 candidate contigs, a contig of 14.8kb showed 81.3% similarity to *Empoasca flavescens* strain As_1 mitochondrion, spanning 14.2kb (Accession No. MK211224.1), with a coverage of 95.9% (E-value = 0.0). All concatenated raw reads were then remapped to the contig showing a mean coverage of 1,689x coverage (**Fig. 1c**).

**Figure 1.**
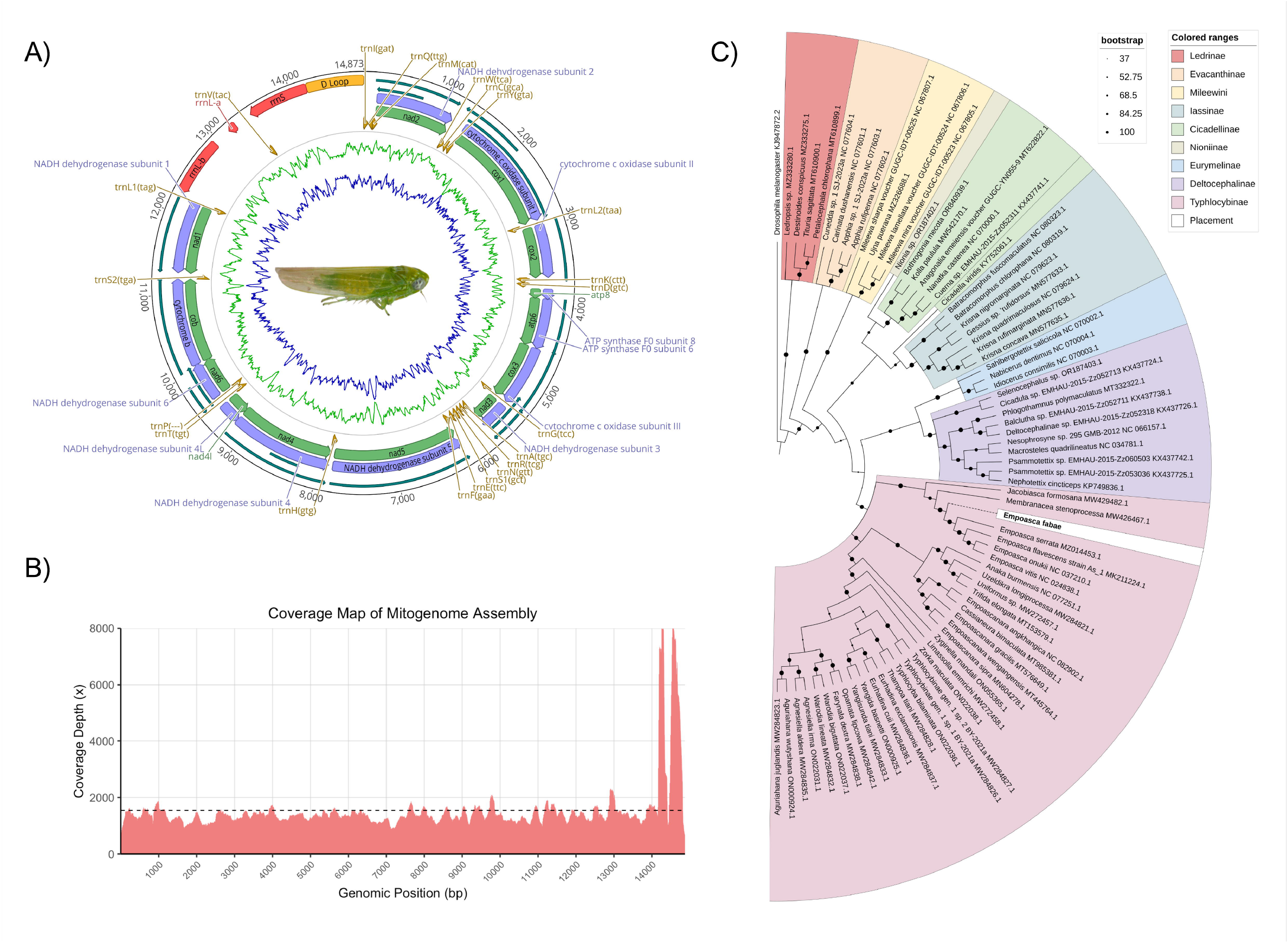
Annotation and coverage map of the *Empoasca fabae* mitogenome, and phylogenetic placement amongst all available unique and complete mitogenomes in NCBI. **A**. *E. fabae* mitogenome annotation showing placement of all 13 PCGs (green) and respective CDSs (purple), rRNAs (red), D Loop (orange), and ORFs (outermost blue lines). **B**. Coverage map of raw sequence reads mapped to the identified mitogenome. **C**. Maximum likelihood phylogenetic tree with bootstrap values and color separated clades to emphasize respective leafhopper subfamilies.

### Annotation

A hybrid annotation approach was used to find and validate the presence of all protein-coding genes (PCGs), ORFs, tRNAs, and rRNAs. Firstly, MITOS2 (Galaxy Version 2.1.9+galaxy0) identified 13 PCGs, 22 tRNAs, two rRNA genes (suspected 16S and 12S) (Bernt et al., 2013). Seqtk (v.1.2) was then used to rearrange the contig based on the cluster of tRNAs before NAD2, specifically trnI, which possesses a GAT anticodon for isoleucine (Shen et al., 2016). Geneious (Dotmatics, USA, v.2024.0) was then used for visualization of the mitogenome (**Fig. 1a**), to plot open reading frames (ORFs) > 300bp, and elucidate base composition (Kearse et al., 2012).

Finally, Mfannot was then used to re-annotate the sequence, confirming 12 of the 13 CDS, with NAD6 being annotated as an ORF covering precisely the same region (Lang et al., 2019). To confirm tRNA predictions, tRNAscan-SE (v.2.0) was used with ‘Infernal’ search mode and without the HMM filter, using invertebrate mitochondrial codes (Lowe & Eddy, 1997). A score cutoff of 10 increased predicted tRNAs from 12 to 14 and lowering the cutoff to 1 increased total tRNA predictions to 19. Each predicted tRNA matched precisely those identified by MITOS2.

### Phylogenetic Analysis

For phylogenetic placement, a multisequence alignment of all 13 PCGs was performed. GenBank was queried using a refined Boolean search to include complete mitochondrial genomes of Cicadellidae subfamilies Aphrodinae, Cicadellinae, Coelidiinae, Deltocephalinae, Eurymelinae, Evacanthinae, Iassinae, Ledrinae, Megophthalminae, Neocoelidiinae, Nioniinae, and Typhlocybinae within the size range of 10,000 to 15,000 bp and published between 2010 and 2024 resulting in 153 species (Benson et al., 2013). All PCGs identified for *E. fabae* were then merged with the queried GenBank file and sorted for each mitochondrial PCG using ‘grep’ and ‘awk’ commands declaring all possible gene ID pattern combinations. Isolated genes were then translated using EMBOSS transeq (v.6.5.7) for their respective amino acid sequences, then aligned using MAFFT (v.7.453) and re-concatenated (Rice et al., 2000, Katoh & Standley, 2013). Genes were then pooled based on their unique accession numbers, with files being omitted from further analysis if they did not contain all 13 PCGs or if sequences were identified as identical duplicates. This resulted in a total of 65 unique species from nine families, including Ledrinae, Mileewini, Cicadellinae, Deltocephalinae, Eurymelinae, Evacanthinae, Iassinae, Nioniinae, and Typhlocybinae, as well as *Drosophila melanogaster* (Accession No. KJ947872.2) as outgroup (**Supplementary Table S1**). The aligned sequences were then analyzed using IQ-TREE (v.2.1.3) using the ModelFinder Plus parameter with limits set restricting model substations to mitochondrial codes (-msub mitochondrial). The consensus tree was then visualized with iTOL (Tree of Life Web Project, USA, v.2024.0) (**Fig. 1c**) (Letunic & Bork, 2021, Nguyen et al., 2013). Branch support was assessed using maximum likelihood method, 1,000 bootstrap replicates and 1,000 approximate likelihood ratio test replicates to ensure robustness.

## RESULTS

The final mitochondrial genome of *E. fabae* was 14,873 bp in length (**Fig. 1a-c**). The genome (Genbank accession no. PQ351619) comprised 13 PCGs, 22 tRNAs, and two rRNAs, with a unique D-loop region (**Table 1**). Total base composition was A (38.8%), T (39.1%), C (11.7%), and G (10.4%). Nearly all PCGs begin with the start codon ATN, consisting of four ATTs, six ATGs, and one ATA. The only two PCGs without ATN codons were NAD5 and COX2, which began with the start codons TTG and GTG, respectively. All predicted PCGs from MITOS2 were confirmed with MFannot, and tRNA predictions were validated using MITOS2 and tRNAscanSE.

**Table 1.**
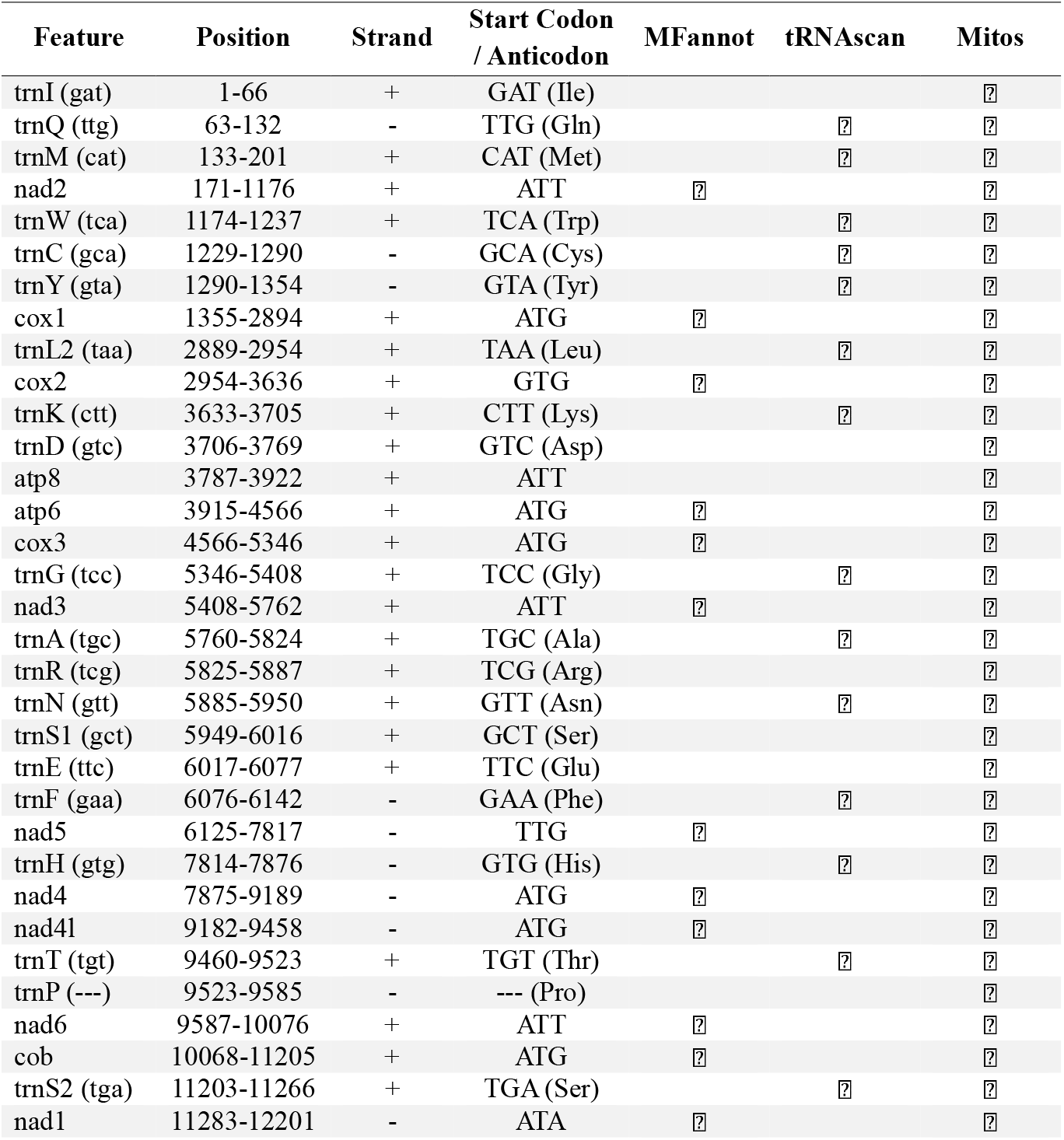
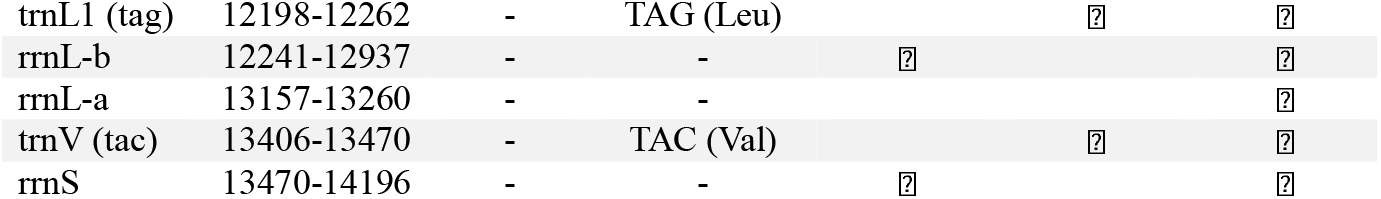
Combined feature table comparing three annotators for the presence of all proteincoding genes and tRNA annotations, including their respective positions and start codons.

After applying the best-fit partitioning scheme to the dataset, the alignment was divided into 13 genes with individual substitution models for each partition, such as the mtInv+F+R6 model for COX1 (**Table 2**). The consensus phylogenetic tree using invertebrate-specific, mitochondrial substation models confirmed the placement of *E. fabae* among leafhoppers within the Typhlocybinae subfamily. Aditionaly, the phylogenetic analysis showed that all members of genus *Empoasca* were clustered together, although a discrete phylogeny of *E. fabae* was shown (**Fig. 1c**).

**Table 2.**
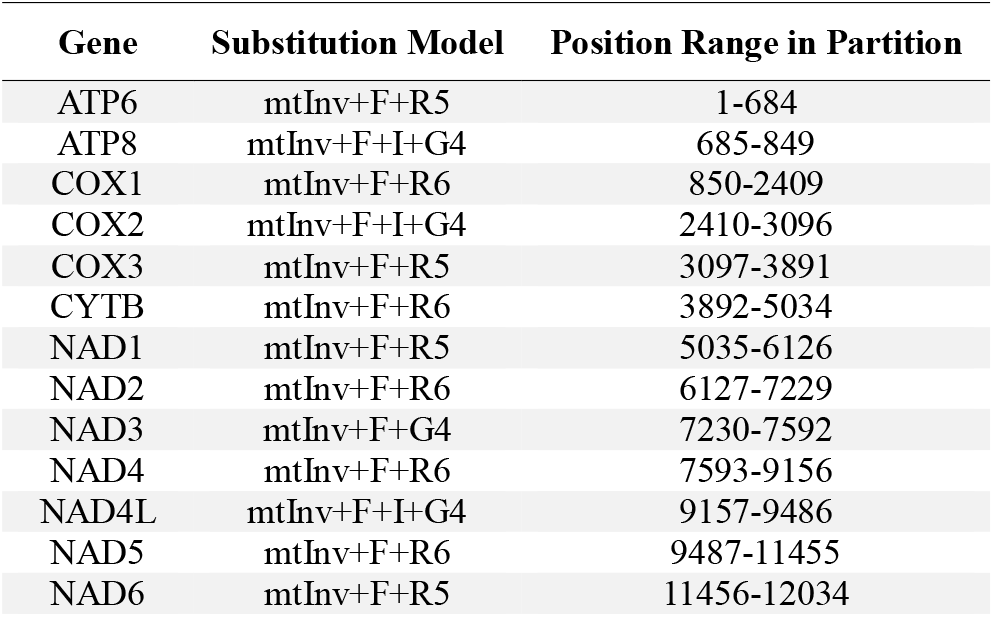
Individual models determined for each protein-coding gene’s amino acid sequence using the IQ-TREE ModelFinder. Models were subsequently used for phylogenetic placement and candidate tree generation.

## DISCUSSION AND CONCLUSIONS

The leafhopper genus *Empoasca* Walsh, 1862 includes 11 subgenera (Oman et al. 1990), with the subgenus *Empoasca* (*Empoasca*) grouping over 600 known species worldwide of which 177 have been reported in North America, 27 in Canada and only *E. bifurcata* DeLong, 1931, and *E. fabae* are present in Quebec (Dmitriev, 2003). Surprisingly, there are only four additional complete mitogenomes available in the GenBank, including those of *E. serrata* Vilbaste, 1965, *E. flavescens* Fabricius, 1794, *E. vitis* Göthe, 1875, and *E. onukii* Matsuda 1952, all produced from specimens collected in China (**Fig. 1c, Supplementary Table S1**).

In this study, we successfully sequenced and annotated the complete mitochondrial genome of *E. fabae* Harris, 1841. This is the first mitogenome for the species, but also the first mitochondrial genome available for a species into the *Empoasca* genus from Canada, North America and the Nearctic region. This study paves the way for more detailed genomic analyses that can improve our understanding of the evolutionary relationships within the *Empoasca* genus and contribute to pest management strategies in agriculture.

## AUTHORS CONTRIBUTION

J.M. Conceptualization, Data curation, Formal analysis, Investigation, Writing - original draft, Writing - review & editing. J.J. Sample collection, Writing - review & editing. S.M. Conceptualization, Writing - review & editing., and E.P.L. Conceptualization, Resources, Supervision, Writing - original draft, Writing review & editing.

## DATA AVAILABILITY STATEMENT

The complete mitochondrial genome assembly data were available in GenBank database under the accession number PQ351619.

## SUPPLEMENTARY FILES

**Supplementary Table S1.**
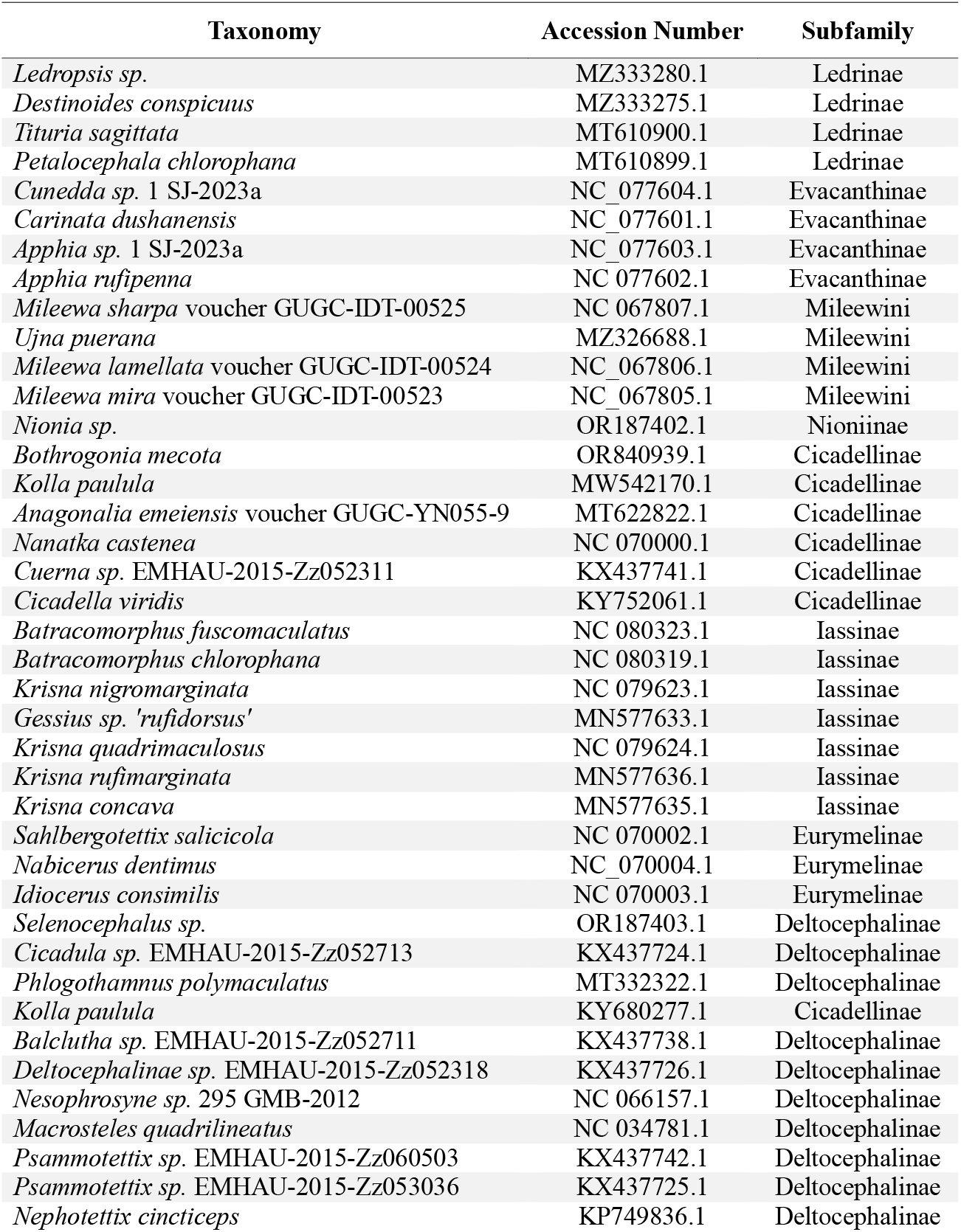
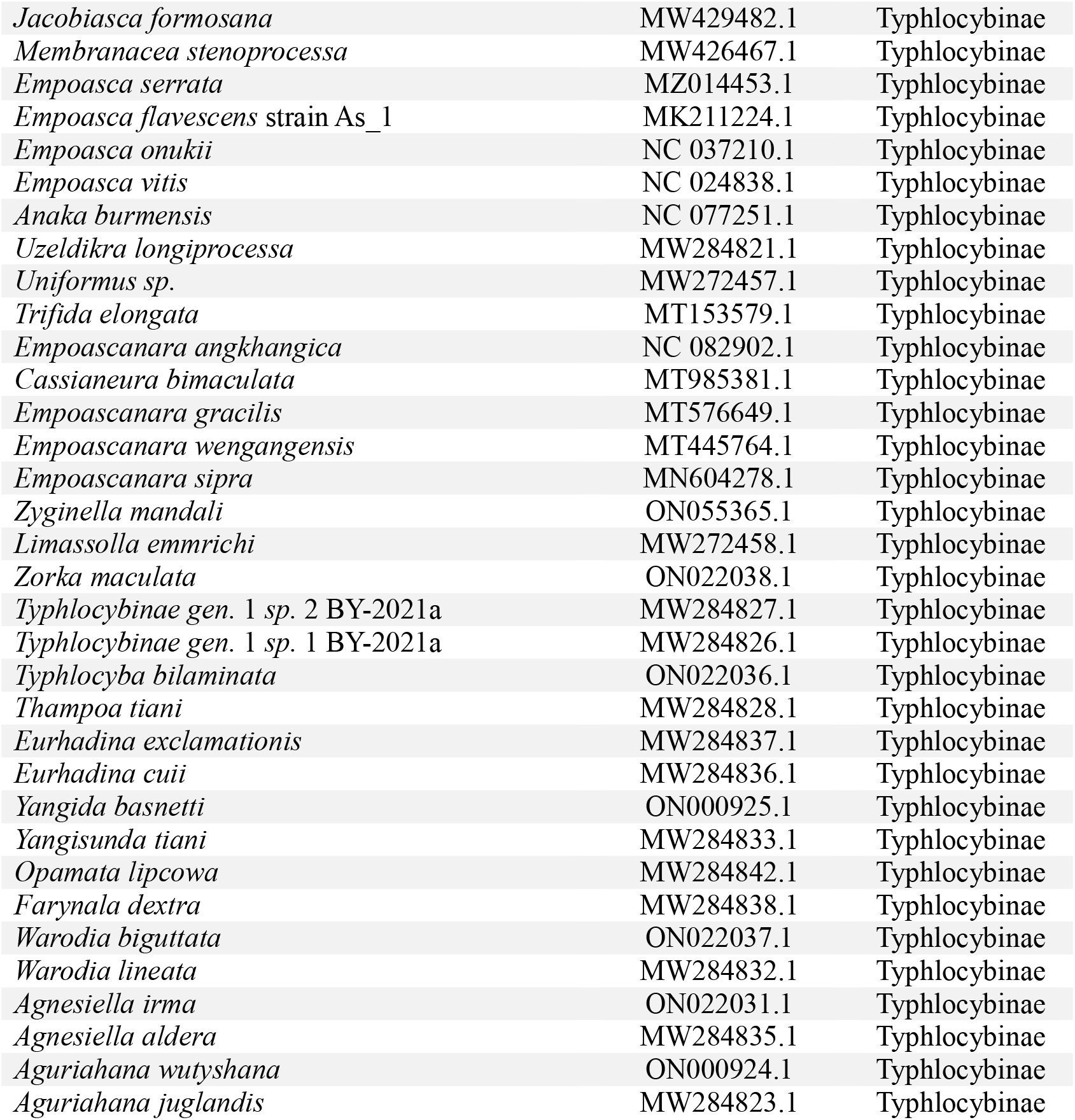
Detailed information of all the mitochondrial genomes used in phylogenetic analysis. All accessions were downloaded from NCBI.

| Taxonomy | Accession Number | Subfamily |
| --- | --- | --- |
| *Ledropsis sp.* | MZ333280.1 | Ledrinae |
| *Destinoides conspicuus* | MZ333275.1 | Ledrinae |
| *Tituria sagittata* | MT610900.1 | Ledrinae |
| *Petalocephala chlorophana* | MT610899.1 | Ledrinae |
| *Cunedda sp.* 1 SJ-2023a | NC_077604.1 | Evacanthinae |
| *Carinata dushanensis* | NC_077601.1 | Evacanthinae |
| *Apphia sp.* 1 SJ-2023a | NC_077603.1 | Evacanthinae |
| *Apphia rufipenna* | NC 077602.1 | Evacanthinae |
| *Mileewa sharpa* voucher GUGC-IDT-00525 | NC 067807.1 | Mileewini |
| *Ujna puerana* | MZ326688.1 | Mileewini |
| *Mileewa lamellata* voucher GUGC-IDT-00524 | NC_067806.1 | Mileewini |
| *Mileewa mira* voucher GUGC-IDT-00523 | NC_067805.1 | Mileewini |
| *Nionia sp.* | OR187402.1 | Nioniinae |
| *Bothrogonia mecota* | OR840939.1 | Cicadellinae |
| *Kolla paulula* | MW542170.1 | Cicadellinae |
| *Anagonalia emeiensis* voucher GUGC-YN055-9 | MT622822.1 | Cicadellinae |
| *Nanatka castenea* | NC 070000.1 | Cicadellinae |
| *Cuerna sp.* EMHAU-2015-Zz052311 | KX437741.1 | Cicadellinae |
| *Cicadella viridis* | KY752061.1 | Cicadellinae |
| *Batracomorphus fuscomaculatus* | NC 080323.1 | Iassinae |
| *Batracomorphus chlorophana* | NC 080319.1 | Iassinae |
| *Krisna nigromarginata* | NC 079623.1 | Iassinae |
| *Gessius sp.* 'rufidorsus' | MN577633.1 | Iassinae |
| *Krisna quadrimaculosus* | NC 079624.1 | Iassinae |
| *Krisna rufimarginata* | MN577636.1 | Iassinae |
| *Krisna concava* | MN577635.1 | Iassinae |
| *Sahlbergotettix salicicola* | NC 070002.1 | Eurymelinae |
| *Nabicerus dentimus* | NC_070004.1 | Eurymelinae |
| *Idiocerus consimilis* | NC 070003.1 | Eurymelinae |
| *Selenocephalus sp.* | OR187403.1 | Deltocephalinae |
| *Cicadula sp.* EMHAU-2015-Zz052713 | KX437724.1 | Deltocephalinae |
| *Phlogothamnus polymaculatus* | MT332322.1 | Deltocephalinae |
| *Kolla paulula* | KY680277.1 | Cicadellinae |
| *Balclutha sp.* EMHAU-2015-Zz052711 | KX437738.1 | Deltocephalinae |
| *Deltocephalinae sp.* EMHAU-2015-Zz052318 | KX437726.1 | Deltocephalinae |
| *Nesophrosyne sp.* 295 GMB-2012 | NC 066157.1 | Deltocephalinae |
| *Macrosteles quadrilineatus* | NC 034781.1 | Deltocephalinae |
| *Psammotettix sp.* EMHAU-2015-Zz060503 | KX437742.1 | Deltocephalinae |
| *Psammotettix sp.* EMHAU-2015-Zz053036 | KX437725.1 | Deltocephalinae |
| *Nephotettix cincticeps* | KP749836.1 | Deltocephalinae |
| *Jacobiasca formosana* | MW429482.1 | Typhlocybinae |
| *Membranacea stenoprocessa* | MW426467.1 | Typhlocybinae |
| *Empoasca serrata* | MZ014453.1 | Typhlocybinae |
| *Empoasca flavescens* strain As_1 | MK211224.1 | Typhlocybinae |
| *Empoasca onukii* | NC 037210.1 | Typhlocybinae |
| *Empoasca vitis* | NC 024838.1 | Typhlocybinae |
| *Anaka burmensis* | NC 077251.1 | Typhlocybinae |
| *Uzeldikra longiprocessa* | MW284821.1 | Typhlocybinae |
| *Uniformus* sp. | MW272457.1 | Typhlocybinae |
| *Trifida elongata* | MT153579.1 | Typhlocybinae |
| *Empoascanara angkhangica* | NC 082902.1 | Typhlocybinae |
| *Cassianeura bimaculata* | MT985381.1 | Typhlocybinae |
| *Empoascanara gracilis* | MT576649.1 | Typhlocybinae |
| *Empoascanara wengangensis* | MT445764.1 | Typhlocybinae |
| *Empoascanara sipra* | MN604278.1 | Typhlocybinae |
| *Zyginella mandali* | ON055365.1 | Typhlocybinae |
| *Limassolla emmrichi* | MW272458.1 | Typhlocybinae |
| *Zorka maculata* | ON022038.1 | Typhlocybinae |
| *Typhlocybinae* gen. 1 sp. 2 BY-2021a | MW284827.1 | Typhlocybinae |
| *Typhlocybinae* gen. 1 sp. 1 BY-2021a | MW284826.1 | Typhlocybinae |
| *Typhlocyba bilaminata* | ON022036.1 | Typhlocybinae |
| *Thampoa tiani* | MW284828.1 | Typhlocybinae |
| *Eurhadina exclamationis* | MW284837.1 | Typhlocybinae |
| *Eurhadina cuii* | MW284836.1 | Typhlocybinae |
| *Yangida basnetti* | ON000925.1 | Typhlocybinae |
| *Yangisunda tiani* | MW284833.1 | Typhlocybinae |
| *Opamata lipcowa* | MW284842.1 | Typhlocybinae |
| *Farynala dextra* | MW284838.1 | Typhlocybinae |
| *Warodia biguttata* | ON022037.1 | Typhlocybinae |
| *Warodia lineata* | MW284832.1 | Typhlocybinae |
| *Agnesiella irma* | ON022031.1 | Typhlocybinae |
| *Agnesiella aldera* | MW284835.1 | Typhlocybinae |
| *Aguriahana wutyshana* | ON000924.1 | Typhlocybinae |
| *Aguriahana juglandis* | MW284823.1 | Typhlocybinae |

